# MetaNet: a scalable and integrated tool for reproducible omics network analysis

**DOI:** 10.1101/2025.06.26.661636

**Authors:** Chen Peng, Zinuo Huang, Xin Wei, Liuyiqi Jiang, Xiaoping Zhu, Zhen Liu, Qiong Chen, Xiaotao Shen, Peng Gao, Chao Jiang

## Abstract

Network analysis is a powerful strategy for uncovering complex relationships in high-throughput omics datasets. However, current tools often lack scalability, flexibility, and native support for multi-omics integration, posing significant barriers for exploring complex biological networks. To address these limitations, we developed MetaNet, a high-performance R package designed to construct, visualize, and analyze biological networks from multi-omics datasets. MetaNet supports highly efficient correlation-based network construction, scalable to datasets with over 10,000 features, and includes extensive layout algorithms and visualization options compatible with both static and interactive platforms. It also provides a comprehensive suite of topological and stability metrics for in-depth network characterization. Benchmarking results show that MetaNet outperforms existing R packages by up to 100-fold in computation time and reduces memory usage by up to 50-fold. We demonstrate its utility through two case studies: (1) a longitudinal analysis of microbial co-occurrence networks showing the dynamics of the airborne microbiome, and (2) an integrative exposome–transcriptome network of more than 40,000 features, uncovering distinct regulatory impacts of biological and chemical exposures. MetaNet bridges the gap between network theory and omics application by offering a robust, reproducible, and biologically informed framework for large-scale, interpretable, and integrative network analyses across diverse omics platforms, advancing systems-level understanding in modern life sciences. MetaNet is available on the Comprehensive R Archive Network (https://cran.r-project.org/web/packages/MetaNet).

## Introduction

Networks, or graphs, are fundamental tools for modeling the intricate relationships in complex biological systems^1^. They provide an abstract yet highly informative representation of interactions between various biological features, ranging from molecular to ecological components^2^. Network theory has profoundly influenced numerous subfields of life sciences, enabling system-level interpretations that transcend individual molecular events. Protein–protein interaction (PPI) networks have elucidated signaling cascades and drug targets^3,4^; gene regulatory networks describe hierarchical control in development and disease^5,6^; metabolic networks map biosynthetic and energy pathways^7,8^; and ecological networks reveal species interactions and community dynamics^9–11^. With the explosion of high-throughput omics technologies, such as metagenomics, transcriptomics, proteomics and metabolomics, network-based approaches have become central to analyzing large-scale, multidimensional biological data^12–15^. These networks help reveal modular structures, infer functional associations, and identify key regulators within and across omics layers^16–18^.

A wide range of tools have been developed for network analysis and visualization. Cytoscape offers a user-friendly platform for visualizing molecular interactions^19^, while Gephi provides efficient layout algorithms for large graphs^20^. The igraph^21^, ggraph^22^, and tidygraph^23^ packages deliver flexible network functions in R and Python; WGCNA is widely used for weighted gene co-expression analysis^24^; and tools like ggClusterNet^25^, microeco^26^, and NetCoMi^27^ extend capabilities for microbiome-specific analyses. Several web-based pipelines (e.g., MENAP^28^, iNAP^29^, and MiCoNE^30^) offer fast, accessible solutions for simple use cases.

However, most existing tools are not designed to meet the full complexity and scale of modern omics data. First, few offer native support for multi-omics integration, limiting their ability to uncover cross-layer associations. Second, computational performance is often a bottleneck—especially for correlation-based network construction on high-dimensional datasets, which may require hours or even days^25^. Third, threshold selection for correlation filtering is typically subjective and arbitrary, potentially biasing network topology and downstream biological interpretation. Fourth, visualization capabilities are often basic, with limited layout flexibility, annotation support, or options for producing publication-quality figures. Finally, many pipelines, particularly web-based tools, lack reproducibility, due to unstable environments, opaque workflows, or the absence of standardized outputs^31^. While such tools remain valuable for general-purpose network analysis, they fall short when applied to the specific demands of omics-scale data: integrative modeling, scalable computation, objective methodology, flexible visualization, and reproducible research.

To address these challenges, we developed MetaNet, a comprehensive and scalable R package tailored specifically for network analysis of omics and multi-omics data. MetaNet fills the above gaps through a unified framework that supports seamless integration of heterogeneous omics layers, enabling the construction of biological networks. By leveraging optimized and parallelized algorithms, it achieves fast correlation-based network construction even on datasets with tens of thousands of features. To improve objectivity, MetaNet incorporates random matrix theory (RMT) for data-driven correlation thresholding, enhancing the reliability of network topology^32,33^. Its extensive visualization module provides over 40 layout algorithms, annotation support, and compatibility with ggplot2, Gephi, and Cytoscape—empowering users to generate customizable, high-quality network figures. MetaNet also emphasizes reproducibility, offering curated example datasets, step-by-step tutorials, and stable version control with seeded randomness. Finally, the package includes a broad suite of topological and stability metrics to support in-depth network interpretation. We demonstrate MetaNet’s capabilities through two representative case studies: a longitudinal microbial co-occurrence network and an integrated exposome–transcriptome network, showcasing its effectiveness in handling complex, large-scale biological data.

## Methods

### Concept Design and Development of MetaNet

MetaNet is an R-based integrative package designed for comprehensive network analysis across diverse omics data, including multi-omics datasets. MetaNet is compatible with operating systems (Windows, MacOS, and Linux) that supports R version 4.0 or higher, and its core functionality is built upon the widely used igraph package. Its architecture comprises several core functional modules: Calculation, Manipulation, Layout, Visualization, Topology analysis, Module analysis, Stability analysis, and I/O (Figure 1A), supporting the end-to-end analytical process from network construction to analysis and visualization. Figure 1B illustrates the main workflow and essential components within MetaNet. The central data structure in MetaNet is the “metanet” object, which extends the widely used “igraph” class from the igraph R package. The “metanet” object is fully compatible with all basic igraph operations and can be converted to a “tbl_graph” object for integration with the ggraph and tidygraph packages. In addition, MetaNet provides a streamlined set of functions tailored to the “metanet” object, facilitating easy construction, annotation, manipulation, visualization, and analysis of biological networks. The workflow is organized through a consistent set of logically structured functions, all beginning with the prefix “c_net_”, making them easy to remember and apply.

**Figure 1.**
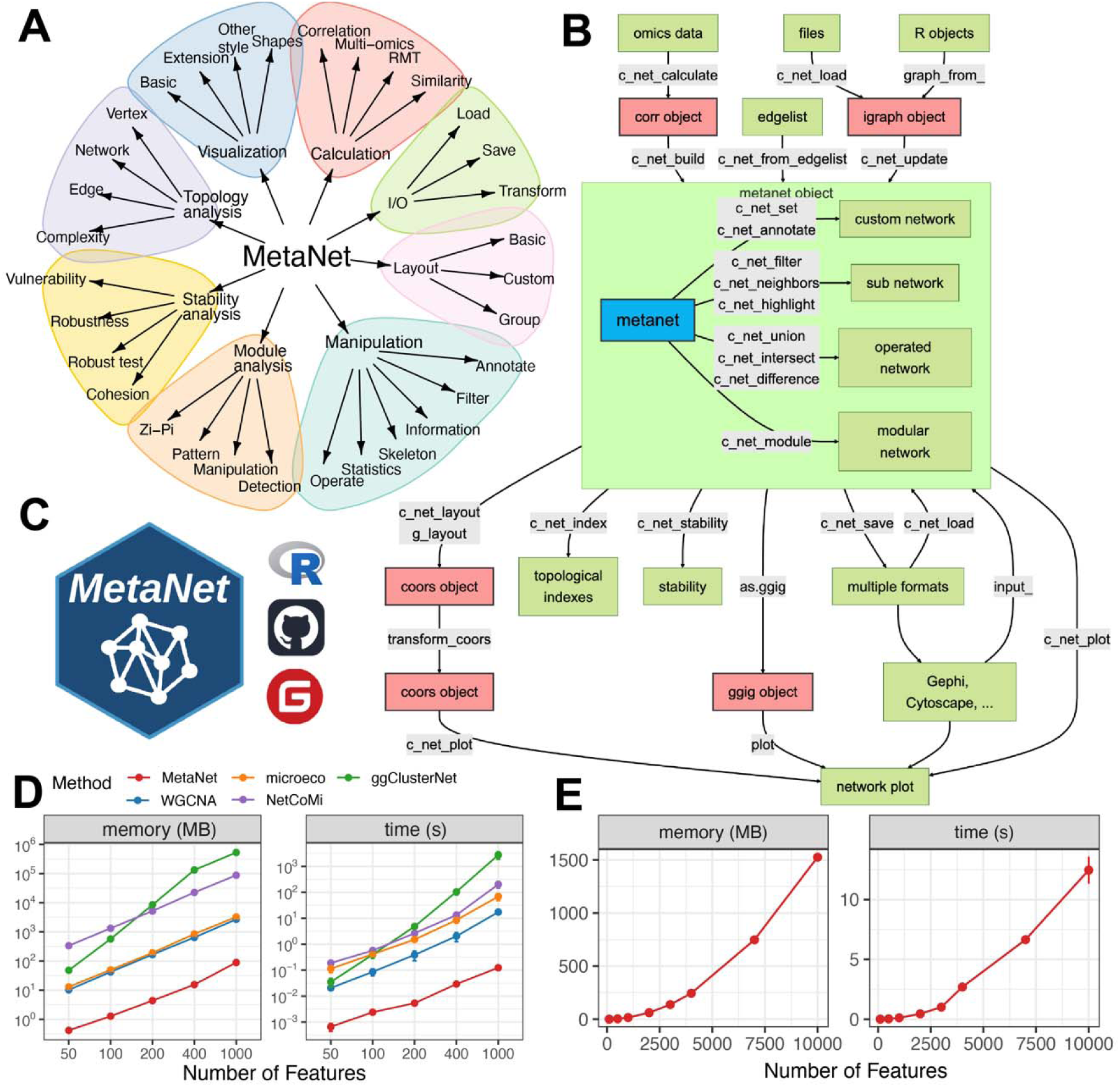
Overview of the MetaNet workflow and its high-efficiency computation. (A) Functional modules of MetaNet, as visualized using MetaNet. (B) Detailed workflow of MetaNet. Green boxes indicate data objects, blue and red boxes represent MetaNet-specific objects, and gray boxes denote core functions. (C) MetaNet logo and its code repositories and platforms. (D) Line plots comparing memory usage and runtime for correlation-based network construction across different R packages. Comparisons were capped at 1000 features because some packages required too many resources and time to process larger networks. Error bars represent standard deviation (SD). (E) Line plots showing MetaNet’s performance on increasingly larger datasets in terms of memory usage and runtime. Error bars represent SD.

#### Data preprocessing

MetaNet provides a broad range of normalization strategies through the “trans” function to accommodate the preprocessing requirements of diverse omics data types (Table S1). For example, transcriptomic data can be transformed using methods such as CPM or log-transformation; microbiome data can be normalized using approaches like aCPM (asinh counts per million) or presence/absence (pa) encoding; and mass spectrometry-based proteomics and metabolomics data can be log1-transformed to reduce skewness and stabilize variance. In addition to normalization, MetaNet also includes utility functions such as “guolv” and “hebing”, which allow users to screen, clean, or combine raw feature tables prior to network construction.

#### Network object construction

Networks can be constructed through several approaches. One method is to calculate and construct networks directly from raw omics data using the “c_net_calculate” and “c_net_build” functions. Alternatively, users may import external network files in formats such as graphml or pajek using “c_net_load” function. Networks can also be built from existing edge list tables using “c_net_from_edgelist”. Finally, “c_net_update” allows for upgrading a conventional igraph object into a fully functional “metanet” object.

#### Annotation of the network

Once a network object is constructed, various operations can be performed on this object. Annotation and attribute assignment are facilitated through “c_net_set” and “c_net_annotate”, enabling advanced downstream data analysis and visualization. And the “get_*” family of functions retrieves tables of network, node, and edge attributes.

#### Manipulation of the network

To extract subnetworks or focus on specific components, functions such as “c_net_filter”, “c_net_neighbors”, and “c_net_highlight” can be used. For comparative analysis across networks, “c_net_union”, “c_net_intersect”, and “c_net_difference” allow for set operations. Community detection or module extraction is achieved using “c_net_module”, which enables users to perform module-based analysis on the resulting network.

#### Layout

For any network, whether original or customized, users can obtain flexible and visually informative layouts using “c_net_layout” and “g_layout”, which provides access to over 40 layout algorithms for generating node coordinates. The “transform_coors” function supports various geometric transformations of layouts, including scaling, aspect ratio adjustments, rotation, mirroring, and pseudo-3D, providing flexible control over network visualization. Visualization of the network is accomplished through the “c_net_plot” function, which offers a variety of parameter settings to help users effectively display network structure and attribute information.

#### Topological analysis

MetaNet also includes advanced functionality for network topology characterization through the “c_net_index” function, which computes 17 widely used topological metrics. Network robustness and structural stability can be evaluated using the “c_net_stability” function, which incorporates multiple stability assessment strategies, particularly relevant for applications such as microbial ecological networks.

#### Accessibility and deployment

MetaNet is completely open source and publicly available on CRAN (https://CRAN.R-project.org/package=MetaNet), GitHub (https://github.com/Asa12138/MetaNet), and Gitee (https://gitee.com/Asa12138/MetaNet). It is actively maintained following CRAN policies (Figure 1C). A comprehensive online manual is also provided to assist users in learning the basics of network analysis and the detailed usage of MetaNet, available at https://bookdown.org/Asa12138/metanet_book/.

### Comparison with MetaNet and other existing tools

To show the computational efficiency, we compared MetaNet with several widely used R packages for biological correlation network construction based on correlation metrics. The “MetaNet::c_net_calculate” function achieves high performance by leveraging vectorized matrix operations with “stats::cor()” and analytically computing p-values using a t-distribution formula, avoiding explicit loops for efficiency. Specifically, we benchmarked the following functions: “MetaNet::c_net_calculate” (v0.2.5), “WGCNA::corAndPvalue” (v1.71), “microeco::trans_network$new” (v0.13.1), “ggClusterNet::corMicro” (v2.00), and “NetCoMi::netConstruct” (v1.1.0). Performance was assessed on a macOS system with an Apple M2 chip. Datasets with varying numbers of features (specifically 50, 100, 200, 400, and 1,000) were used to evaluate memory usage and computation time for each method. These metrics were measured using the “bench::mark” function (v1.1.2), and each test was repeated 20 times to ensure reliability. Statistical comparisons of memory usage and computation time at each feature number were performed using the Wilcoxon rank-sum test, revealing significant improvements of MetaNet over other R packages (p < 0.001).

### Case study

We utilized a recently published longitudinal multi-omics dataset that included transcriptomic profiles, biological exposome (microbial exposome), and chemical exposome profiles, collected from individuals in an underwater environment^34^. To study species interaction networks, we first selected microbial taxa that appeared in at least 2 out of 24 samples across 6 long-term sampled individuals, corresponding to a prevalence threshold of more than 10%. This filtering resulted in a total of 914 microbial species retained for network construction. Species co-occurrence networks were built using Spearman correlation analysis. Only edges with an absolute correlation coefficient (|ρ|) greater than 0.6 and BH-adjusted p-values less than 0.05 were retained, consistent with thresholds used in prior microbial ecological studies^35^. To evaluate the modularity structure and determine the number of modules in each network, we employed a fast greedy modularity optimization algorithm^36^. Each module represents a set of taxa with stronger intra-group than inter-group connectivity, reflecting potential ecological or functional coherence. To capture temporal variation in microbial interactions, we extracted sample-specific subnetworks of bacteria based on species presence (defined as abundance > 0 in a given sample). For subnetwork of each sample, we calculated a suite of topological indices to characterize the structure of microbial communities over time. These metrics included edge density, proportion of negative edges, average degree (connectivity), average path length, network diameter, clustering coefficient (transitivity), eigenvector centrality, betweenness centrality, closeness centrality, degree centrality, and natural connectivity.

For the integrative multi-omics network analysis, correlation networks were constructed between exposure layers (biological and chemical) and the transcriptome using Spearman correlation. For each pair of omics datasets, correlation matrices were computed by calculating Spearman’s rank correlation coefficients and associated p-values across all features. To retain only robust associations consistent with the previous study, variable pairs with |ρ| > 0.6 (for chemical–transcriptome pairs) or > 0.5 (for biological–transcriptome pairs) and BH-adjusted p-values < 5e–4 were included in the final networks. Finally, for genes significantly associated with either microbial or chemical exposures, we conducted functional enrichment analysis using over-representation analysis (ORA) against the KEGG^37^ and Gene Ontology^38^ (GO) databases by ReporterScore (v0.2.2) package^39^. This allowed us to identify molecular pathways and biological processes most affected by the different types of environmental exposures.

## Results

### Efficient and scalable network computation enables analysis of larger omics datasets in MetaNet

Network analysis has become a cornerstone in many omics disciplines. Before constructing networks, different omics data types—including microbiome, transcriptome, proteome, and metabolome—require appropriate preprocessing to ensure data quality and reliability. MetaNet offers a broad range of normalization strategies (Table S1, see Methods), enabling effective preprocessing across omics types.

Network construction begins with computing pairwise relationships using statistical strategies^40^. The primary method includes similarity or correlation-based approaches such as “Spearman”, “Pearson”, and “Bray-Curtis”. These methods generate similarity matrices between features, followed by randomization-based significance testing and multiple testing correction to retain only statistically meaningful associations. Users can choose from several widely used correction methods, including the Benjamini–Hochberg false discovery rate (FDR), Bonferroni, or Holm adjustment, all of which are implemented using “c_net_calculate “.

Pairwise correlation computation is central to most network-based omics tools. However, with the rapidly increasing scale of omics datasets, many existing tools struggle with high computational demands. MetaNet addresses this limitation through optimized vectorized matrix algorithms for calculating correlation coefficients and corresponding p-values, significantly reducing memory use and runtime (Figure 1D, see Methods). Benchmarking showed that MetaNet completed correlation-based analysis on datasets with fewer than 1,000 features in under 0.2 seconds and using less than 100 MB of memory (Figure 1D). This performance surpassed other tools by 100- to 10,000-fold in speed (p < 0.001). While other tools may take over an hour on large datasets, MetaNet maintained low resource usage, with both memory and time scaling approximately quadratically with feature number (Figure 1E). These efficiencies make MetaNet well-suited for high-throughput network construction across large omics datasets.

Correlation-based association networks are widely adopted due to their simplicity and robustness to noise, but a key limitation is the subjective selection of thresholds. Most studies rely on manually defined criteria (e.g., |r| > 0.6 and p < 0.05) to define edges of networks^35^, introducing inconsistencies and potential biases. To address this, MetaNet incorporates RMT, a statistically grounded method for identifying optimal thresholds^32,33^. RMT automatically determines a correlation cutoff that minimizes spurious edges based on the structure of the data. This integration provides users with a data-driven approach to define the r_threshold parameter for network construction (Figure S1A and S1B). While MetaNet primarily supports correlation-based methods, it is also compatible with the result of alternative approaches to network inference. These include mutual information-based methods, which capture non-linear relationships, and partial correlation-based approaches, which control for indirect associations.

### Streamlined functional tools for network annotation, manipulation, and comparison in MetaNet

Once a network is constructed, exploratory analysis and flexible manipulation are essential to extract biological insights. MetaNet provides a suite of functions to support these tasks. For instance, the “get_*” family of functions retrieves tables of network, node, and edge attributes, enabling inspection and statistical summarization. The resulting “metanet” object is fully compatible with all basic igraph operations and can also be converted to a “tbl_graph” object for seamless integration with the ggraph and tidygraph packages.

In omics and multi-omics studies, networks are often annotated with external data such as abundance profiles, taxonomy, or clinical metadata. The “c_net_set” function attaches multiple annotation tables to a network object and automatically configures internal properties for network visualization (Figure 2B). This includes setting color schemes, line types, node shapes, and generating appropriate legends.

**Figure 2.**
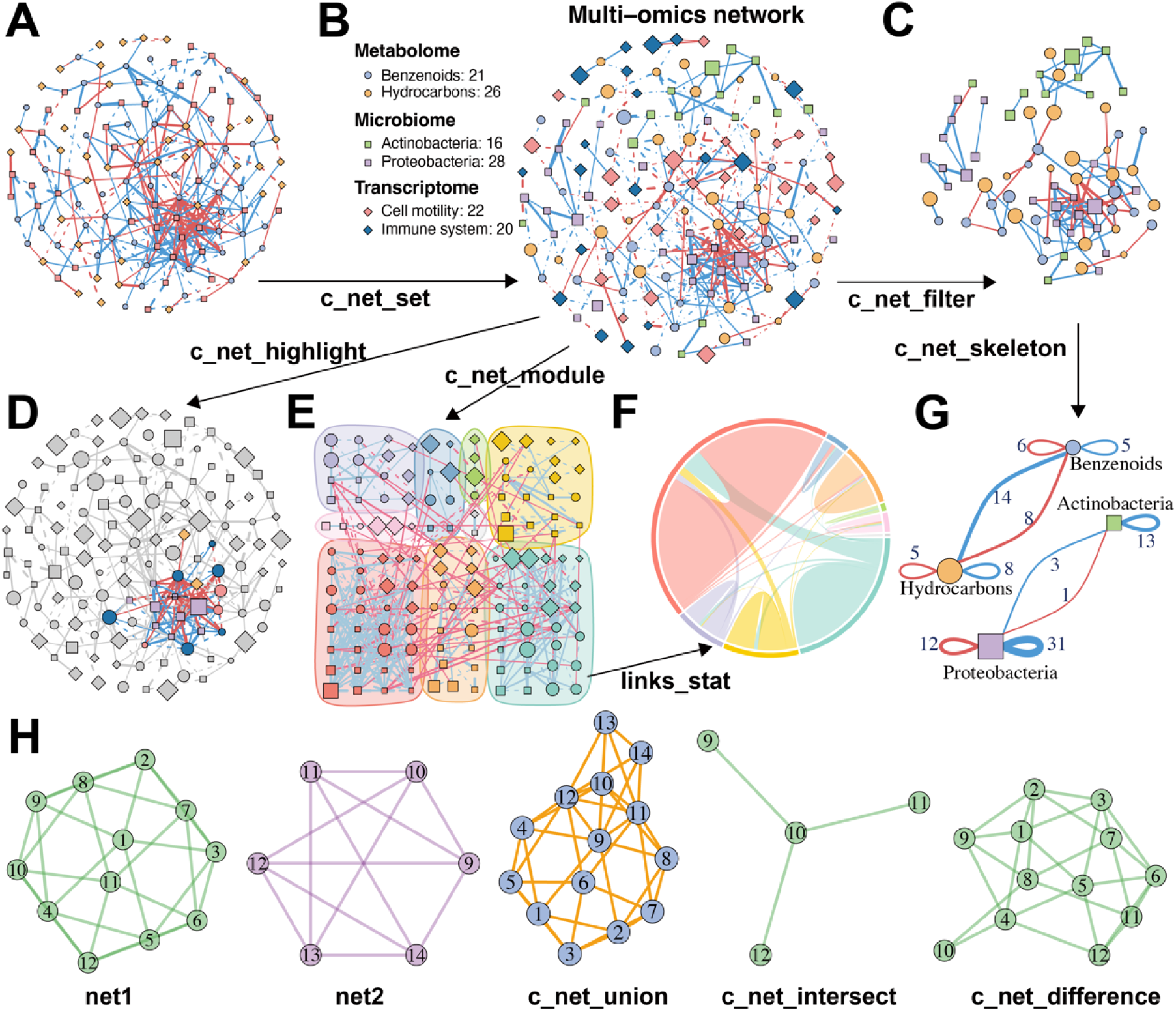
MetaNet supports flexible and intuitive network manipulation. (A) Initial multi-omics network constructed without annotations. (B) Annotated multi-omics network using the “c_net_set” function. Node shape indicates the types of omics data, color represents the subtypes of omics data, size denotes average abundance, edge color indicates positive or negative correlation, edge type (solid or dashed) distinguishes intra- and inter-omics connections, and edge width reflects the absolute value of the correlation coefficient. (C) Subnetwork filtered from intra-omics interactions between the Microbiome and Metabolome layers using “c_net_filter”. (D) Highlighted nodes centered on “Dongia_mobilis” and its neighbors using “c_net_highlight”. (E) Community detection and modular visualization using “c_net_module”. (F) Chord diagram displaying the proportion of edges between modules. (G) Skeleton network across omics subtypes at a grouped level using “c_net_skeleton”. (H) Operations among networks: “c_net_union” merges net1 and net2, “c_net_intersect” extracts shared nodes and edges, and “c_net_difference” isolates net1-specific nodes and edges. All networks shown are based on simulated data and are for illustrative purposes only.

After annotation and visual customization, researchers may need to focus their analysis on specific network regions. This is particularly common in multi-omics integration scenarios. The “c_net_filter” function enables the extraction of sub-networks using flexible and combinable filters (Figure 2C). To visually emphasize specific regions of interest within a network, the “c_net_highlight” function emphasizes specific nodes or edges (Figure 2D).

In complex networks, modules or communities refer to subgraphs with densely connected nodes^41^. These structures often represent biologically meaningful groups, such as functionally related gene clusters or regulatory units. MetaNet supports module detection with the “c_net_module” function, which includes multiple community detection algorithms (Figure 2E). The resulting modules can be visualized using chord or Sankey diagrams, which clearly depict the proportions and inter-module connections (Figure 2F). To further support analysis at the group level, the “c_net_skeleton” function enables statistical summarization of edge origins and targets across group levels, enhancing interpretability in multi-condition or longitudinal datasets (Figure 2G).

Comparative analysis across multiple networks is critical in omics research. For example, researchers may identify differential edges between experimental groups or track stable subnetworks during transitions. MetaNet facilitates such comparisons through functions that compute intersections, unions, and differences between multiple networks (Figure 2H), offering a flexible framework for network-based comparisons and evolutionary analysis.

### Advanced network layout and visualization support in MetaNet

Layout is a critical component of network visualization, as a well-designed layout can significantly enhance the interpretability of network structures^42^. MetaNet stores layout coordinates in a flexible object called “coors”, which allows users to control, reuse, and transfer layout settings with ease. The “c_net_layout” function provides access to over 40 layout algorithms for generating node coordinates (Figure 3A), including several new layouts as well as adaptations from the igraph^21^ and ggraph^22^ packages. In addition to conventional layouts, MetaNet introduces the “spatstat_layout” method, which enables users to constrain layout generation within a user-defined polygon or along its edges. This layout function supports uniform or random node distributions inside custom shapes. For example, arranging a network within a star (Figure 3B) or mapping it to a geographic region like Australia (Figure 3C). MetaNet also offers convenient interoperability with interactive visualization platforms such as Gephi and Cytoscape. Users can easily import layouts that were computed or manually adjusted in these external tools.

**Figure 3.**
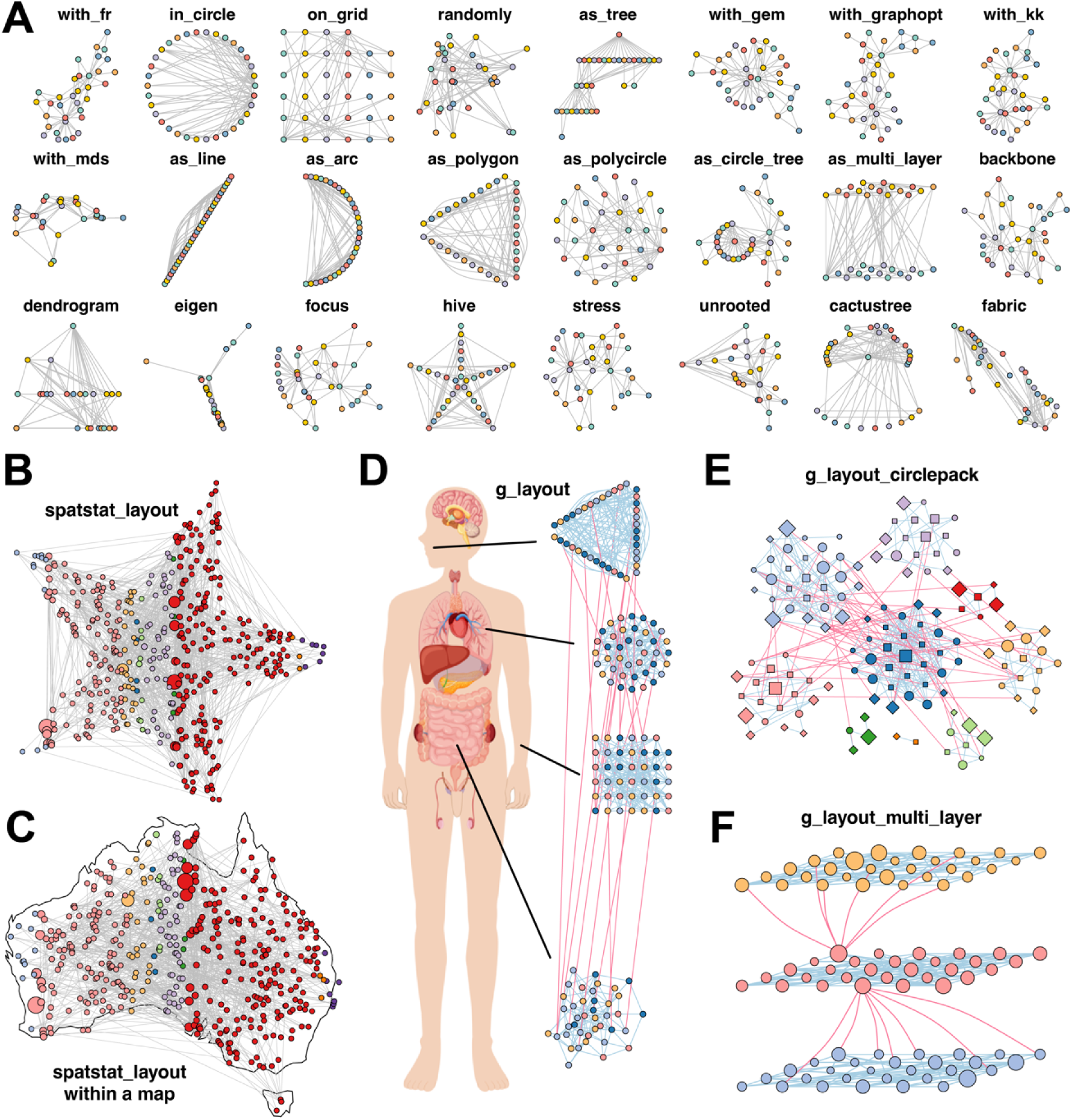
MetaNet enables diverse and powerful network layout strategies. (A) The application of 24 out of more than 40 built-in layout algorithms from “c_net_layout” on the Zachary Karate Club network was provided by the igraph package. (B) Layout generated within a star using “spatstat_layout”. (C) Layout generated within the map of Australia using “spatstat_layout”. (D) Grouped network layout consisting of four subgroups arranged with “with_fr()”, “on_grid()”, “as_polycircle(3)”, and “as_polygon(3)” within a human-body schematic. All visualization elements were rendered with MetaNet without manual adjustments. (E) Modular network visualized using “g_layout_circlepack”. (F) A three-layer modular structure visualized using “g_layout_multi_layer”. All networks shown are based on simulated data.

For networks with grouping variables, MetaNet provides an advanced layout interface via the “g_layout” function. It lets users define spatial configurations for each group, including positioning, scaling, and internal layout strategies. Multiple layout types can be combined in a single visualization. The “g_layout” function returns a “coors” object, which can be nested or recombined with subsequent “g_layout” calls to create highly customized multi-level layouts. For example, a co-abundance network of human microbiomes from multiple body sites can be efficiently arranged using a single call to “g_layout” (Figure 3D). This compound layout strategy is also particularly useful for displaying modular structures within networks. For instance, “g_layout_circlepack” can be used to clearly illustrate the spatial distribution of detected modules in a compact circular packing layout (Figure 3E), while “g_layout_multi_layer” introduces a multi-layer pseudo-3D representation that highlights inter-modular relationships (Figure 3F). Other group layouts are shown in Figure S2.

MetaNet’s “c_net_plot function” offers extensive parameters for visual customization (Table S2), enabling precise control over nodes, edges, modules, and legends to create high-quality visualizations. By default, MetaNet uses R’s base plotting system inherited from igraph. However, users preferring ggplot2 can use the “as.ggig” function to convert networks into ggplot-compatible structures. This makes it possible to apply advanced ggplot2 functions such as “labs”, “theme”, and “ggsave” to refine figures (Figure S3A). MetaNet also supports exporting visual content to external tools such as NetworkD3, Gephi, and Cytoscape for extended visualization workflows (Figure S3B-D).

### Extended support for specialized and database-linked biological networks

MetaNet provides native support for a variety of specialized network types frequently used in bioinformatics workflows, enabling researchers to visualize and explore biological relationships beyond conventional correlation or interaction networks.

MetaNet allows the construction of Venn-style networks to illustrate set relationships across sample groups. These provide a more informative alternative to traditional Venn diagrams by displaying explicit connections and network structure (Figure 4A). Tree-structured data, such as taxonomies or gene ontology hierarchies, can be visualized using the built-in “as_circle_tree” layout, offering a clear and compact representation of hierarchical relationships (Figure 4B). MetaNet further supports pie-node visualization, where each node encodes multivariate annotations, such as group-specific abundances. This approach allows compositional data to be embedded directly in the network structure (Figure 4C).

**Figure 4.**
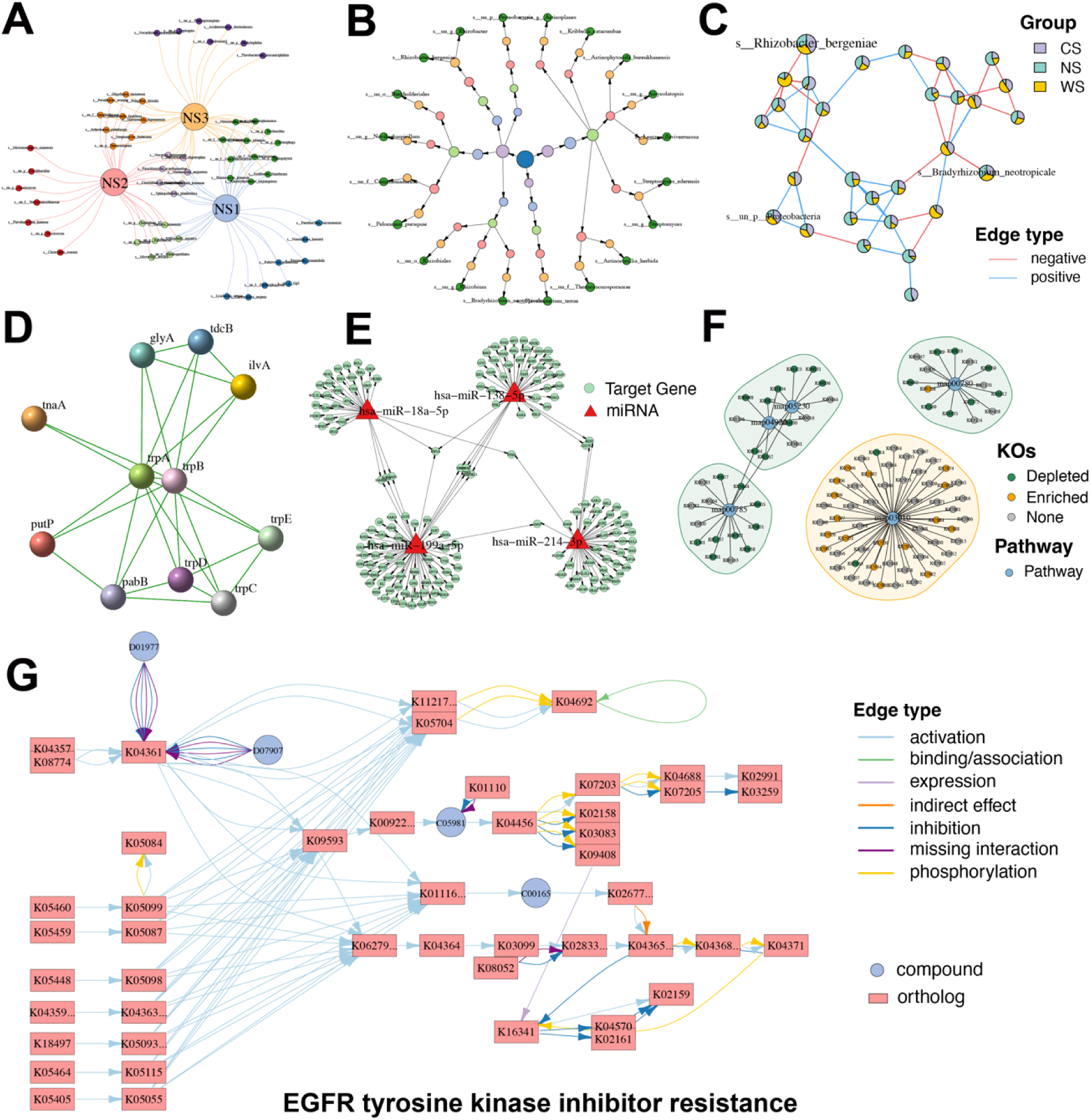
Diverse specialized network visualizations by MetaNet. (A) Venn-style network: Large nodes represent groups, while smaller nodes denote individual elements within each group, enabling visualization of shared and unique components. (B) Hierarchical tree network: Nodes are organized based on classification hierarchy. Node color corresponds to the taxonomic or categorical level. (C) Pie-node network: Each node is displayed as a pie chart, where slice colors indicate relative abundance across different groups. (D) Protein–protein interaction (PPI) network: Extracted from the STRING database, showing experimentally validated and predicted molecular interactions among proteins. (E) miRNA–gene regulatory network: Sourced from the miRTarBase database, illustrating experimentally supported regulatory relationships between miRNAs and their target genes. (F) KEGG KO–pathway association network: The network shows KEGG orthologs (KOs) involved in selected biological pathways. Small nodes represent KOs, and large nodes represent pathways. KO nodes are colored by their expression trend. Shaded regions surrounding pathways indicate whether the pathway is globally up-regulated (orange) or down-regulated (green). (G) KEGG pathway-specific network: Network representation of the “EGFR tyrosine kinase inhibitor resistance” pathway. Rectangular nodes denote KEGG orthologs, circular nodes indicate compounds, and edge colors reflect interaction types.

Beyond generic network types, MetaNet is compatible with biological networks from external databases. For example, protein–protein interaction (PPI) networks obtained from the STRING database^43^ can be imported and visualized with customized layout and annotations (Figure 4D). Similarly, miRNA–target gene regulatory networks from miRTarBase^44^, which are experimentally validated, can be represented to explore post-transcriptional regulatory mechanisms (Figure 4E).

MetaNet also integrates with the ReporterScore, an R package we previously developed for functional enrichment analysis^39^. Using the results of pathway enrichment, users can directly visualize relationships between KEGG orthologs (KOs) and their associated pathways (Figure 4F). Furthermore, MetaNet supports direct rendering of any KEGG pathway map through a specified pathway ID, enabling fully annotated and modifiable visualizations (Figure 4G).

Together, these extended features highlight MetaNet’s versatility in accommodating diverse biological network types, integrating with external knowledge bases, and enhancing the interpretability of complex multi-omics analyses.

### Comprehensive network topology and stability analysis in MetaNet

Network topology refers to the structural patterns formed by the connections between nodes and edges, reflecting both global architecture and local importance in complex biological systems^45^. In omics research, topological analysis is crucial for understanding molecular interactions and functional organization. MetaNet offers a broad suite of topological metrics for characterizing networks at global and local levels. Global metrics describe the overall network structure, including density, average degree (connectivity), average clustering coefficient, average path length, natural connectivity, and others. (Table S3). These metrics quantify essential properties of biological networks such as redundancy, robustness, and signal propagation potential. For instance, average path length reflects the typical number of steps required to traverse the network between nodes, which can be interpreted as an estimate of signaling efficiency in metabolic or gene regulatory networks^46^. Local metrics evaluate the importance or centrality of individual nodes or edges (Table S4), helping identify key regulators or bottlenecks within the system.

To assess structural significance, MetaNet can generate random networks using the Erdős–Rényi model^47^ with identical numbers of nodes and edges as the observed network (Figure S4A). This allows for comparison with real omics data-derived networks, which often exhibit scale-free, small-world, modular, and hierarchical features^17^. The “fit_power” function tests for scale-freeness by fitting a power-law to the degree distribution (Figure S4B), while “smallworldness” computes the small-world index σ. Modular structure is a hallmark of biological networks, representing clusters of closely connected nodes often corresponding to functionally related features (Figure S4C). MetaNet implements multiple community detection algorithms via the “c_net_module” function, allowing users to examine expression or abundance patterns within modules (Figure S4D). Furthermore, based on the Zi-Pi method^48^, MetaNet also classifies nodes into four topological roles: peripherals, connectors, module hubs, and network hubs, offering functional insights into individual nodes (Figure S4E and S4F).

Beyond topology, network stability is essential in modeling robustness in molecular systems, ecosystems, and metabolic regulation^16^. MetaNet incorporates several algorithms to assess structural and ecological stability, which can be computed using “parallel::detectCores” for enhanced efficiency. For the structural robustness test, MetaNet calculates natural connectivity as nodes are progressively removed from the network^49^ (Figure S5A). The rate of decline in connectivity reflects the network’s resilience to perturbation^50^. Robustness is assessed by simulating node removals and tracking survival based on the abundance-weighted mean interaction strength^51^ (Figure S5B). Vulnerability reflects a node’s contribution to global efficiency, indicating its critical role in network communication^51^ (Figure S5C). Cohesion indices, both positive and negative, measure cooperation and competition within microbial communities^52^ (Figure S5D and S5E).

### Case 1. Longitudinal dynamics of a microbial co-occurrence network

To demonstrate the flexibility of MetaNet in diverse and integrated omics analyses, we applied it to a recently published individual-level longitudinal study involving multi-omics data^34^. In this study, the research team developed wearable passive samplers to perform high-resolution temporal profiling of both chemical and biological exposomes in 19 individuals exposed to a specialized environment. The data included integrated transcriptome and exposome profiles, offering a unique opportunity to examine the effects of environmental perturbations on individual health. Here, we focused on the microbial exposome component, representing the airborne microbiota encountered by each participant over time. Timepoint A represented baseline conditions in a natural environment, while timepoints B to D were recorded in the exposure environment.

We first constructed a global microbial co-occurrence network (Figure 5A), which included 871 microbial species spanning four taxonomic kingdoms (Figure 5B). Using a greedy modularity optimization algorithm^36^, we identified six distinct modules with varying intra-module species compositions (Figure 5A). The degree distribution of this network followed a power-law distribution, indicating scale-free properties (Figure 5C). This suggests that the observed network possesses characteristics of a complex system. Within each module, we analyzed microbial abundance patterns across timepoints. For example, members of Module M3 showed a consistent decline in relative abundance over time. Topological role classification using the Zi-Pi method revealed 13 module hubs and 19 connectors, likely critical for network integrity and inter-module communication.

**Figure 5.**
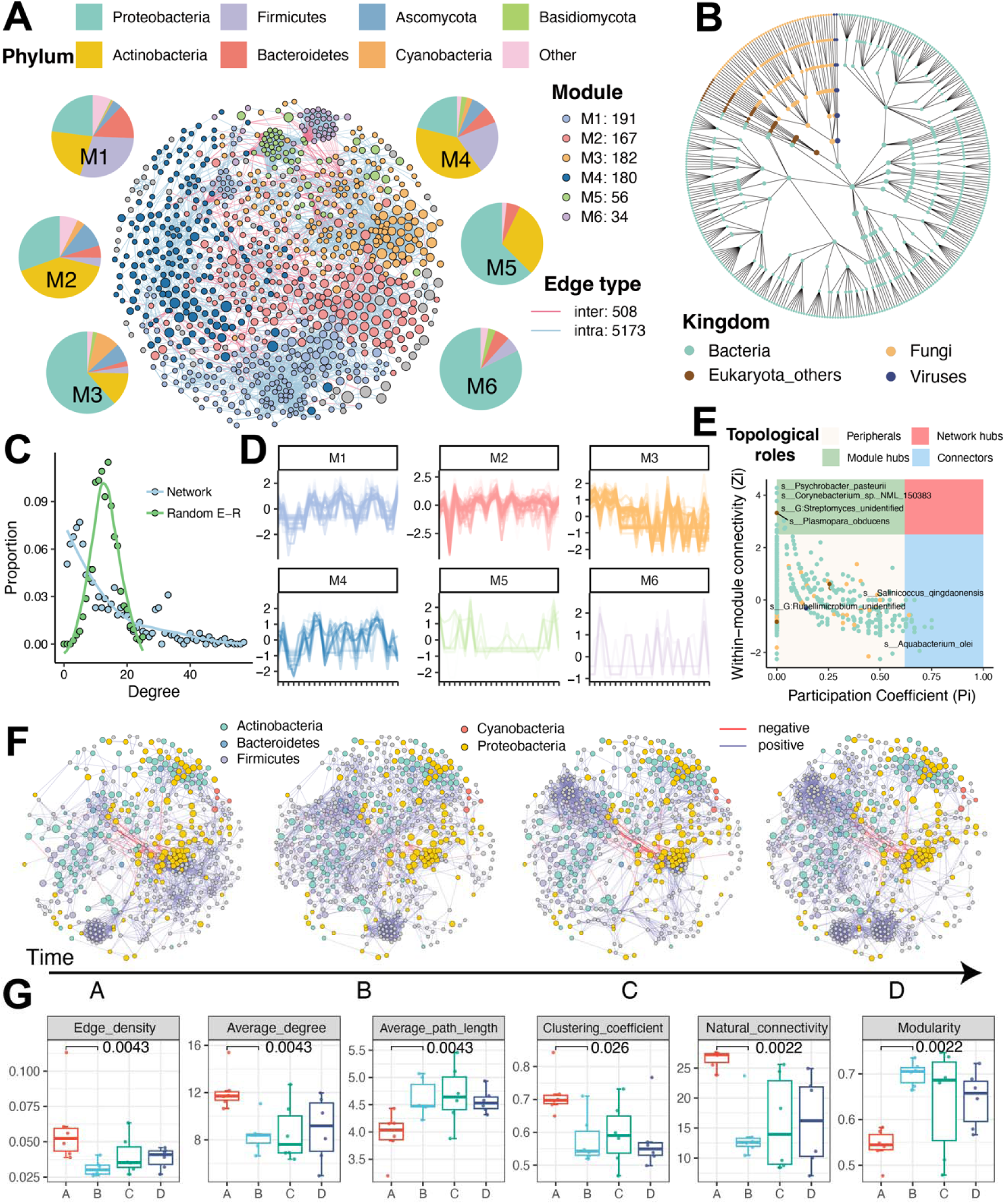
Modularity and temporal dynamics of the microbial co-occurrence network. (A) Species-level microbial co-occurrence network constructed from all microbial exposure samples, showing six modules (M1 to M6). Node color indicates module membership, node size reflects relative abundance, and edge color distinguishes intra-versus inter-module connections. (B) Phylogenetic relationship network of all species in panel A, arranged using the “as_circle_tree” layout. (C) Comparison of degree distribution between the empirical network in panel A and a randomized network with the same number of nodes and edges. (D) Temporal abundance profiles of species within each module. The y-axis represents the scaled abundance of species, while the x-axis represents individual samples sorted by time point (E) Key microbial taxa identified based on topological role classification using the Zi-Pi framework. (F) Subnetwork dynamics across four exposure stages. Node color represents bacterial phylum, node size reflects relative abundance, and gray nodes denote non-core species with presence or abundance changes over time. (G) Changes in global network topological metrics across different stages. P-values for comparisons between timepoints A and B were calculated using the Wilcoxon rank-sum test.

We also extracted sub-networks of bacteria for each exposure timepoint (Figure 5F). A subset of microbial species was found to change in presence or abundance over time (colored in gray). Further topological analysis indicated major shifts from timepoint A to B (Figure 5G). Compared to the pre-exposure at timepoint A, networks at timepoint B to D exhibited increased modularity and average path length, accompanied by decreases in global efficiency, clustering coefficient, and natural connectivity. Biologically, these alterations may reflect community fragmentation and a breakdown of previously stable bacterial interactions. Increased modularity indicates the emergence of isolated subcommunities with limited cross-talk, while reduced clustering and efficiency suggest that bacterial communication and co-functionality became impaired. Together, these patterns point to a loss of ecological resilience, consistent with the notion that the airborne microbiome was destabilized and functionally weakened under the specialized exposure conditions^35^.

### Case 2. Multi-omics integrated network reveals differential impacts of biological and chemical exposome on the transcriptome

We extended our analysis of the longitudinal multi-omics dataset by performing integrated network analysis between the exposome (both biological and chemical) and the host transcriptome. This analysis aimed to uncover the temporal associations between the environmental exposome and gene expression. We used MetaNet to efficiently compute correlation networks between 35,587 transcriptomic genes, 2,955 microbial species, and 3,729 chemical exposome features (Figure 6A). Our results revealed that 590 microbial taxa were significantly associated with 1,983 genes, with the majority of edges representing positive correlations. In contrast, 245 chemical exposures showed significant associations with 1,026 genes, among which negative correlations were more prevalent. To visualize the most prominent associations, we presented a multi-omics integrative network based on the strongest correlations (Figure 6B). The network topology showed a clear distinction: microbial exposures were predominantly positively correlated with gene expression, while chemical exposures showed a bias toward negative associations (Figure 6C). Among microbial taxa, *Microbacterium lacticum* and *Aureobasidium melanogenum* exhibited the highest number of significant associations with host genes. For chemical compounds, the top contributors included (SR)- or (Rs)-4-methyl-2,3-pentanediol, indole, α,3-dichlorotoluene, and 3-ethylcatechol, with the first compound showing the highest number of gene associations.

**Figure 6.**
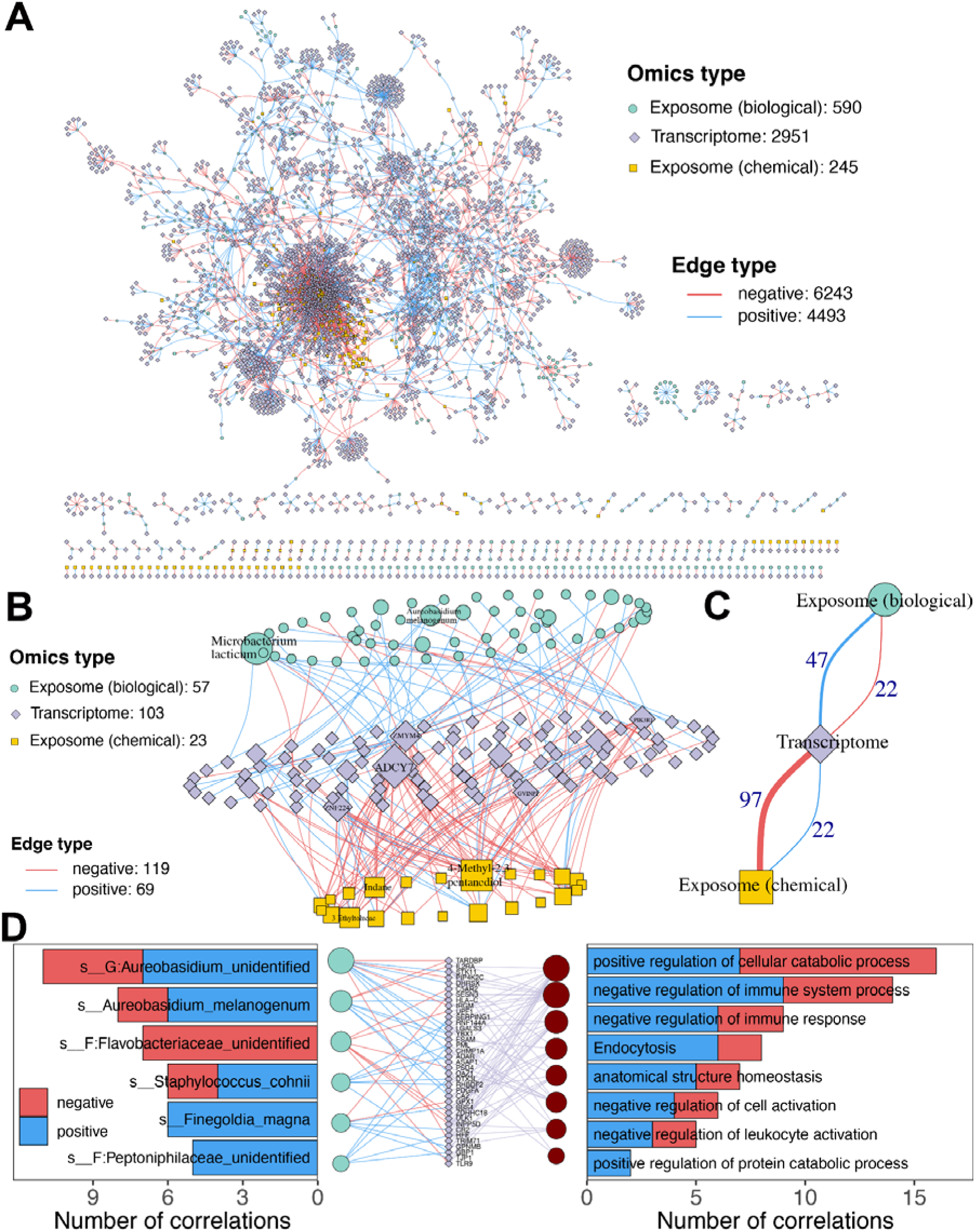
Integrated Network Analysis of Exposome–Transcriptome Interactions. (A) Spearman correlation-based multi-omics network linking all microbial and chemical exposures to transcriptomic data. (B) Spearman correlation-based multi-omics network showing the most prominent associations. Links with |ρ| > 0.7 (for chemical–transcriptome pairs) or > 0.6 (for biological–transcriptome pairs) are included. Only the top 10 ranked node labels are shown. Node size reflects degree centrality. (C) Skeleton structure of the network in panel B, highlighting the core architecture of microbial and chemical associations with genes. (D) Network representations of significantly correlated genes and enriched pathways for biological exposures. Bar charts on either side indicate the number of positively and negatively correlated connections for each exposure.

We further mapped the genes correlated with either microbial or chemical exposures to the GO and KEGG databases to identify enriched biological functions and pathways^53^. Notably, eight pathways significantly associated with the microbial exposome were identified, most of which were related to the negative regulation of immune responses (Figure 6D). Among these, several gram-positive anaerobic bacteria emerged as potential immunological stressors due to their positive correlations with genes such as *ADAR*, *GBP1*, and *RHBDF2*, which are key regulators of immune signaling^54^. *Streptococcus salivarius*, known for its immunosuppressive properties, showed a significant positive correlation with *HMGB1*, a central mediator in inflammatory signaling, suggesting potential cross-talk between exposure and host inflammatory responses^55^.

In contrast, the chemical exposome was predominantly linked to disease-related pathways (Figure S6). These included neurodegenerative diseases such as Parkinson’s disease and Alzheimer’s disease, as well as pathways involved in DNA damage response and cellular stress. Of particular concern, compounds such as benzene, ethylbenzene, and xylene—known to cross the blood–brain barrier—have been implicated in neuropsychiatric disorders, including attention deficits and cognitive decline^56^. Polycyclic aromatic hydrocarbons (PAHs), another class of chemicals identified in the dataset, have been associated with elevated risks of DNA damage and carcinogenesis^57^.

In summary, this case study demonstrates how MetaNet can facilitate integrated multi-omics network analysis. It highlights distinct molecular signatures associated with biological versus chemical exposures and underscores how exposomes may influence disease-related pathways and health outcomes through specific gene–environment interactions.

## Discussion

In this study, we present MetaNet, a scalable, flexible, and biologically-informed R package for network analysis of omics and multi-omics data. By integrating network construction, visualization, topological analysis, and cross-layer integration into a single reproducible workflow, MetaNet addresses key limitations present in existing tools. Its ability to handle thousands of features, support diverse network types, and generate high-quality, customizable visualizations makes it particularly suited for modern systems biology. Through two case studies, we demonstrate MetaNet’s practical value in extracting biologically meaningful insights from complex datasets. Benchmark comparisons confirm its computational advantages, while modular function design ensures extensibility and user adaptability.

There is a growing ecosystem of network analysis tools, each with unique strengths and design philosophies^58^. Cytoscape offers excellent visualization capabilities and plugins, Gephi supports large-scale dynamic network layouts, igraph provides high-performance network operations in R and Python, WGCNA excels at weighted gene co-expression analysis, and packages like NetCoMi and ggClusterNet enable specialized microbial and ecological network construction. The diversity of available platforms allows researchers to choose methods tailored to specific tasks^59^.

With advances in high-throughput technologies, omics datasets are becoming increasingly complex. Modern assays capture tens of thousands of features with higher resolution, studies now span multiple omics layers, and sample sizes continue to grow in cohort-scale and longitudinal designs. These trends call for analytical tools that are both scalable and versatile. MetaNet contributes to this landscape by offering several indispensable features: ultra-fast computation of correlation-based networks, seamless integration of multi-omics layers, comprehensive support for biological network types, flexible and customizable visualizations, and advanced metrics for topological and stability analysis. It is particularly well-suited for microbial ecological networks, multi-omics interaction networks, and studies requiring precise, reproducible plotting—such as repeated network rendering with fixed coordinate systems. In addition, MetaNet supports the visualization and analysis of specialized biological networks (e.g., KEGG pathways, PPI, regulatory networks) and continues to expand its capabilities. Fully open-source and designed for extensibility, MetaNet functions both as a standalone platform and as a flexible foundation for building future multi-omics network pipelines.

Although MetaNet offers substantial advancements in omics-scale network analysis, certain limitations remain. The current optimization primarily focuses on correlation-based networks, which, though widely used, may overlook complex, non-linear, or indirect associations. This highlights the need to integrate broader inference strategies, including partial correlations^60^, mutual information^61^, or regression-based approaches. As omics datasets continue to grow in size and complexity, there is also increasing demand for more efficient algorithms—not only for network construction, but also for advanced layout computation and topological analysis at scale. Continuous optimization and architectural refinement will be required to meet these demands. Moreover, MetaNet faces broader challenges common in biological network analysis^1^, including the lack of standardized protocols for network construction, such as similarity metrics, thresholding, and null model selection^62^. Frequently used topological metrics— such as node degree, connectivity, or clustering coefficient—are often intercorrelated and can lead to contradictory biological interpretations, complicating downstream analysis and biological inference^16^.

Looking forward, MetaNet’s architecture provides fertile ground for future expansion. First, incorporating additional methods for network inference, such as regression-based, Bayesian, or machine learning approaches, could improve the robustness and interpretability of constructed networks. Second, deeper integration of curated biological knowledge from databases like STRING and KEGG could enable hybrid networks that combine empirical and prior data. Third, as single-cell omics becomes increasingly prevalent, supporting sparse matrix representations and network construction tailored to single-cell data would further extend MetaNet’s applicability. Finally, expanding support for ecological networks, such as food webs or host–parasite systems, along with corresponding ecological metrics (e.g., trophic level, connectance), would open new avenues for microbial ecology and environmental omics studies.

Taken together, MetaNet offers a robust solution for multi-omics network analysis and is poised to grow into a central platform for the field through ongoing development and community engagement.

## Data availability

Code is available as an open-source R package ‘MetaNet’, which can be downloaded from GitHub (https://github.com/Asa12138/MetaNet). The main analysis scripts (Rmarkdown format) and source data are available from GitHub (https://github.com/Asa12138/Analysis_code/tree/main/MetaNet_figures).

## Acknowledgments

We are grateful to our colleagues at the core facility of the Life Sciences Institute, especially the NECHO high-performance computing cluster. This research was partly supported by grants from NSFC 82341109 and 82173645 and the Fundamental Research Funds for the Central Universities.

## Author contributions

C.P. and C.J. conceived the study. C.P. developed the R package. C.P., Z.H., X.W., and L.J. collected all datasets. C.P. completed the main benchmarking and case study analyses. Z.H., X.W., L.J., X.Z., Z.L., Q.C., X.S. and P.G. contributed to the analyses. C.P. and C.J. drafted and revised the manuscript with input from other authors.

## Supplementary material

**Figure S1.**
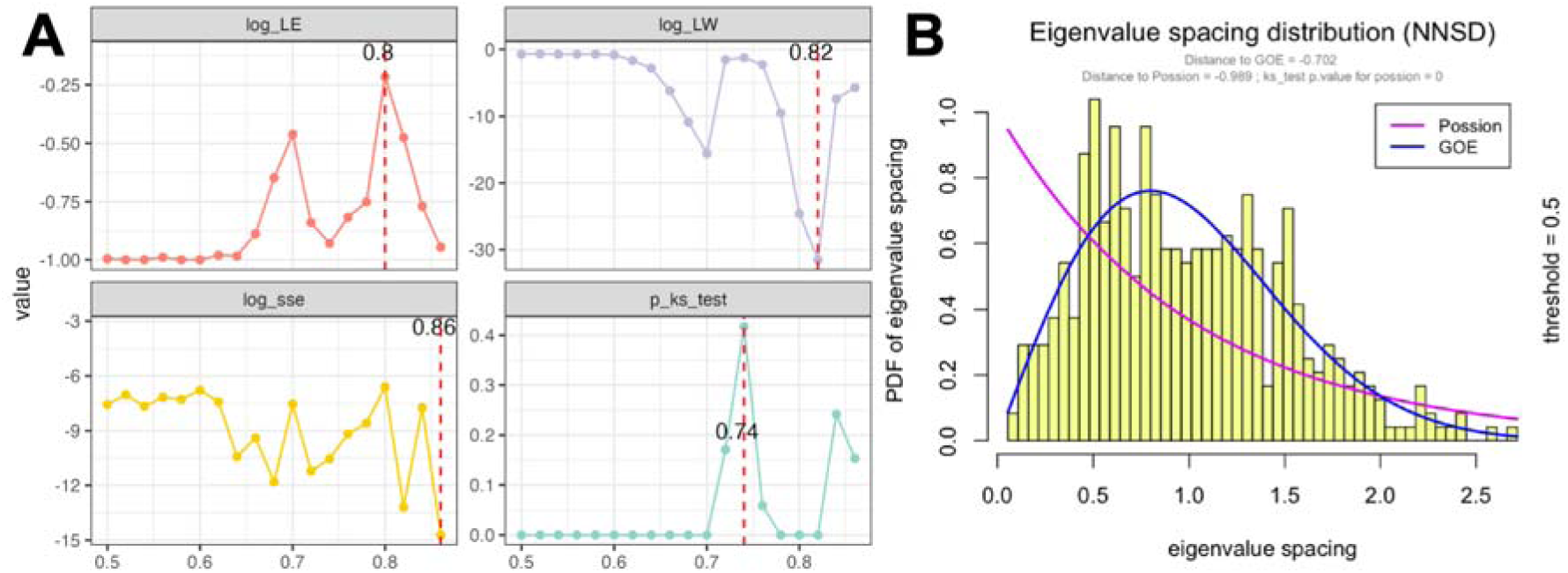
Evaluation of RMT-based network threshold optimization. (A) Line plots showing changes in key random matrix theory (RMT)-based statistics under different correlation thresholds. A meaningful threshold is indicated by higher values of log_LE and p_ks_test, and lower values of log_LW and log_SEE. These metrics help identify the optimal “r_threshold” for robust network construction. (B) Probability density function (PDF) of eigenvalue spacing distribution when the correlation threshold is set to 0.5.

**Figure S2.**
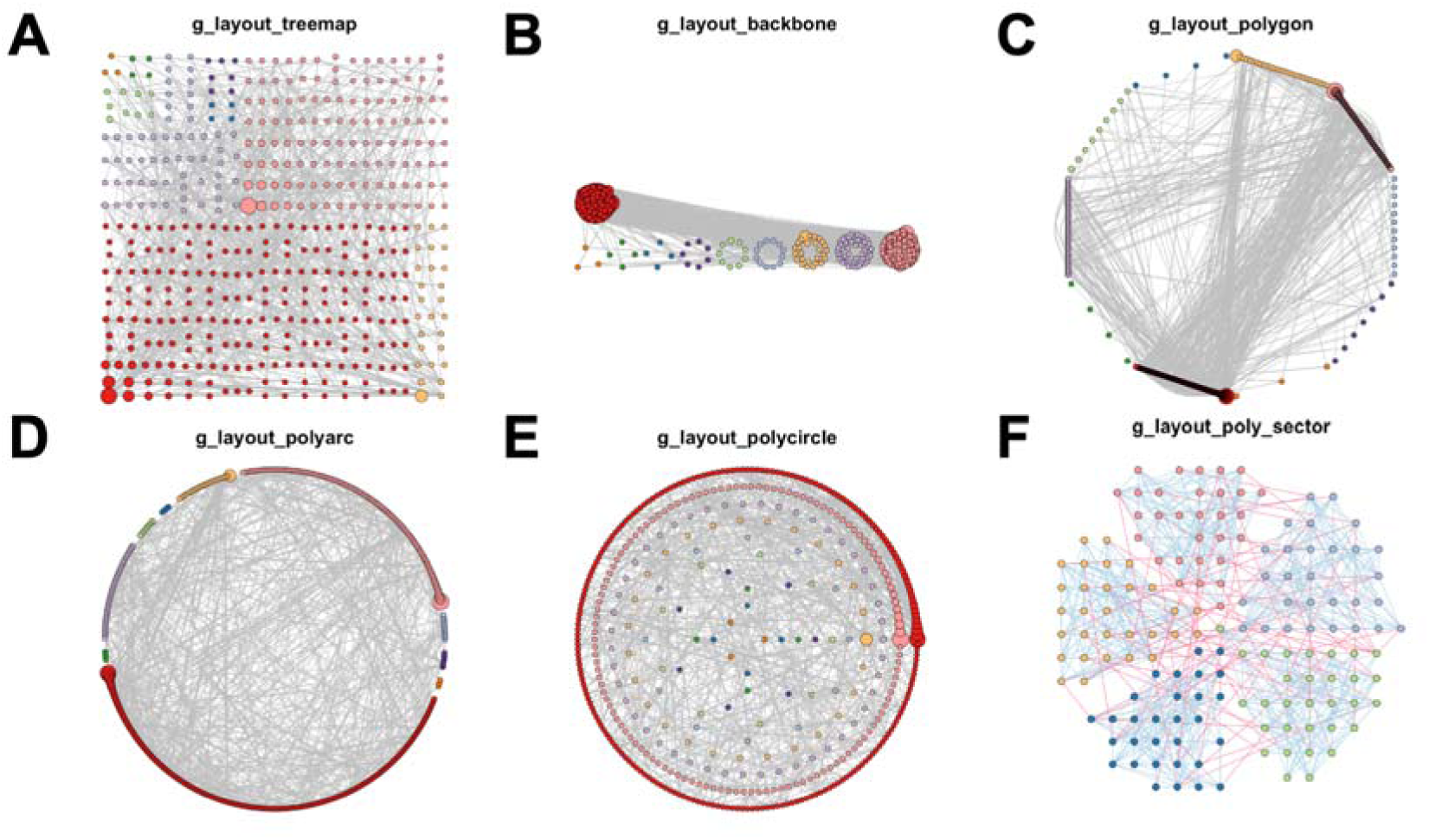
Preset layout options for group-based network visualization. (A) g_layout_treemap, (B) g_layout_backbone, (C) g_layout_polygon, (D) g_layout_polyarc (E) g_layout_polycircle, and (F) g_layout_poly_sector.

**Figure S3.**
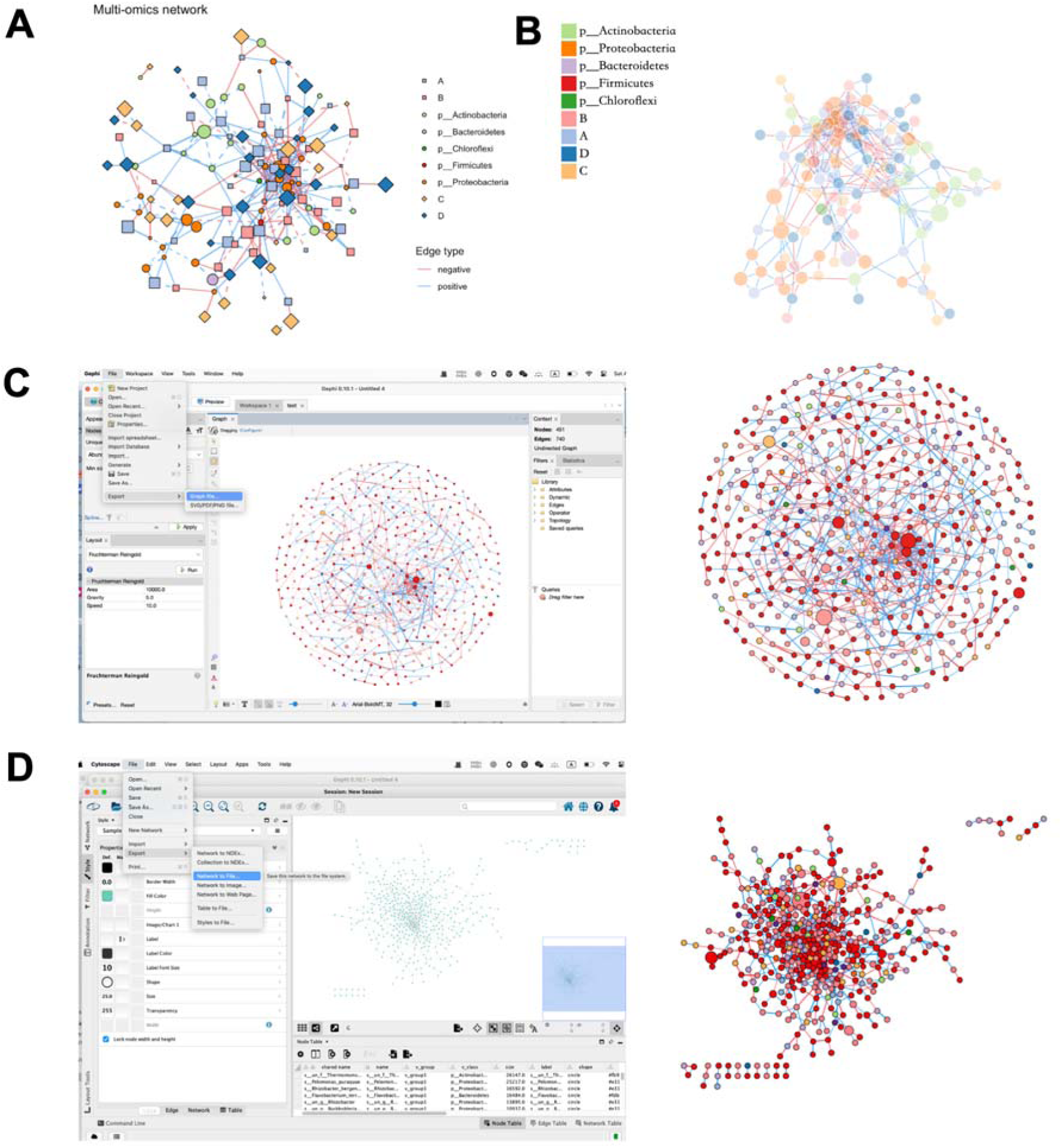
MetaNet compatibility with external visualization tools. Extension of MetaNet to multiple external visualization and analysis platforms, including: (A) ggplot2-based rendering via “as.ggig”, (B) D3.js-based interactive networks via “ netD3plot”, (C) exported static layout rendering in Gephi, and (D) import and annotation within Cytoscape.

**Figure S4.**
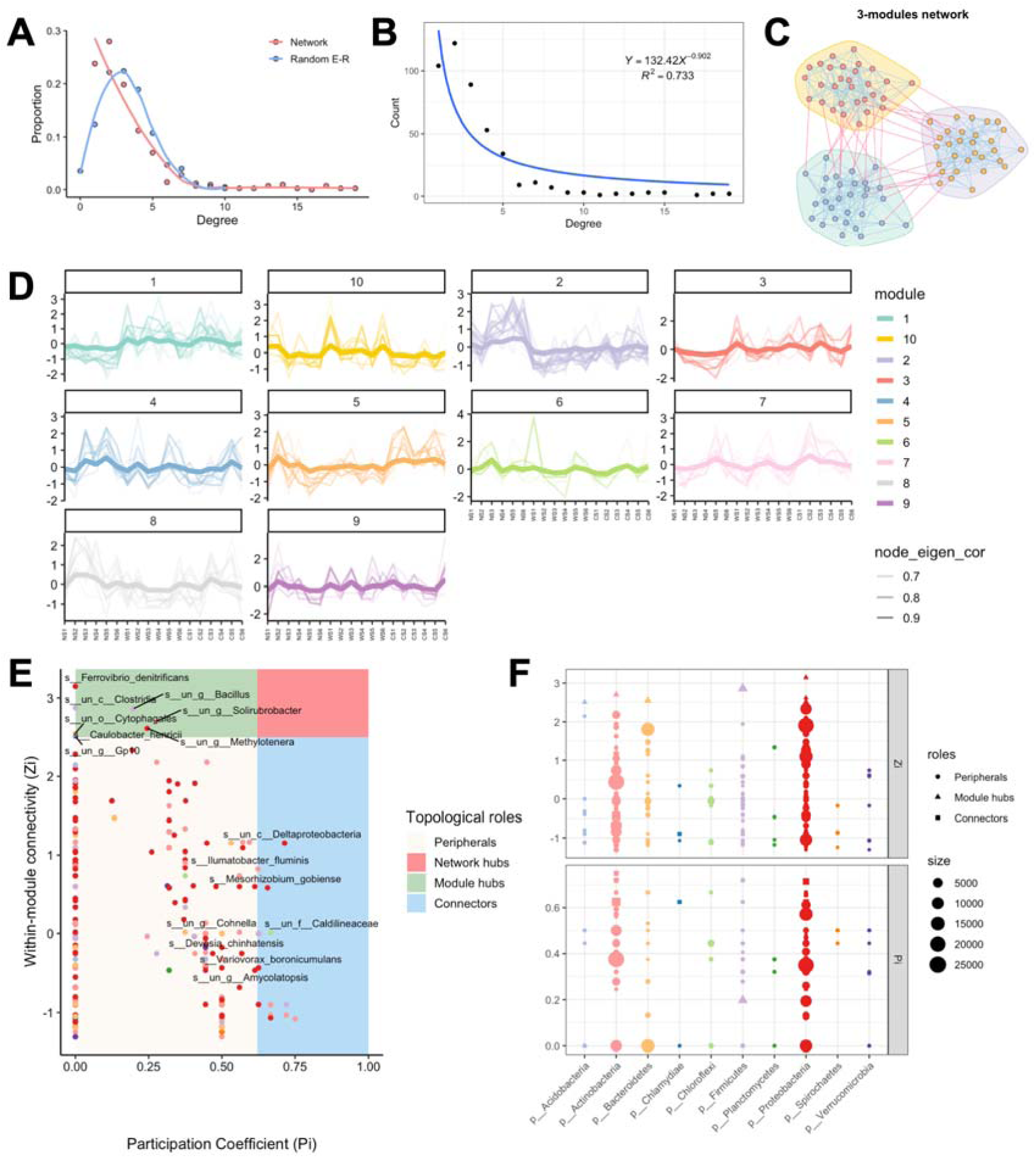
Structural properties and modular analysis of microbial networks. (A) Degree distribution comparison between the constructed microbial network and a random network with the same number of nodes and edges. (B) Power-law fitting of degree distribution in the constructed network. (C) Example of a module-level subnetwork extracted from the global network. (D) Line plots of intra-module species abundance across samples. (E) Classification of node topological roles using participation coefficient (P) and within-module connectivity (Zi). (F) Distribution of Zi and P values across different microbial phyla.

**Figure S5.**
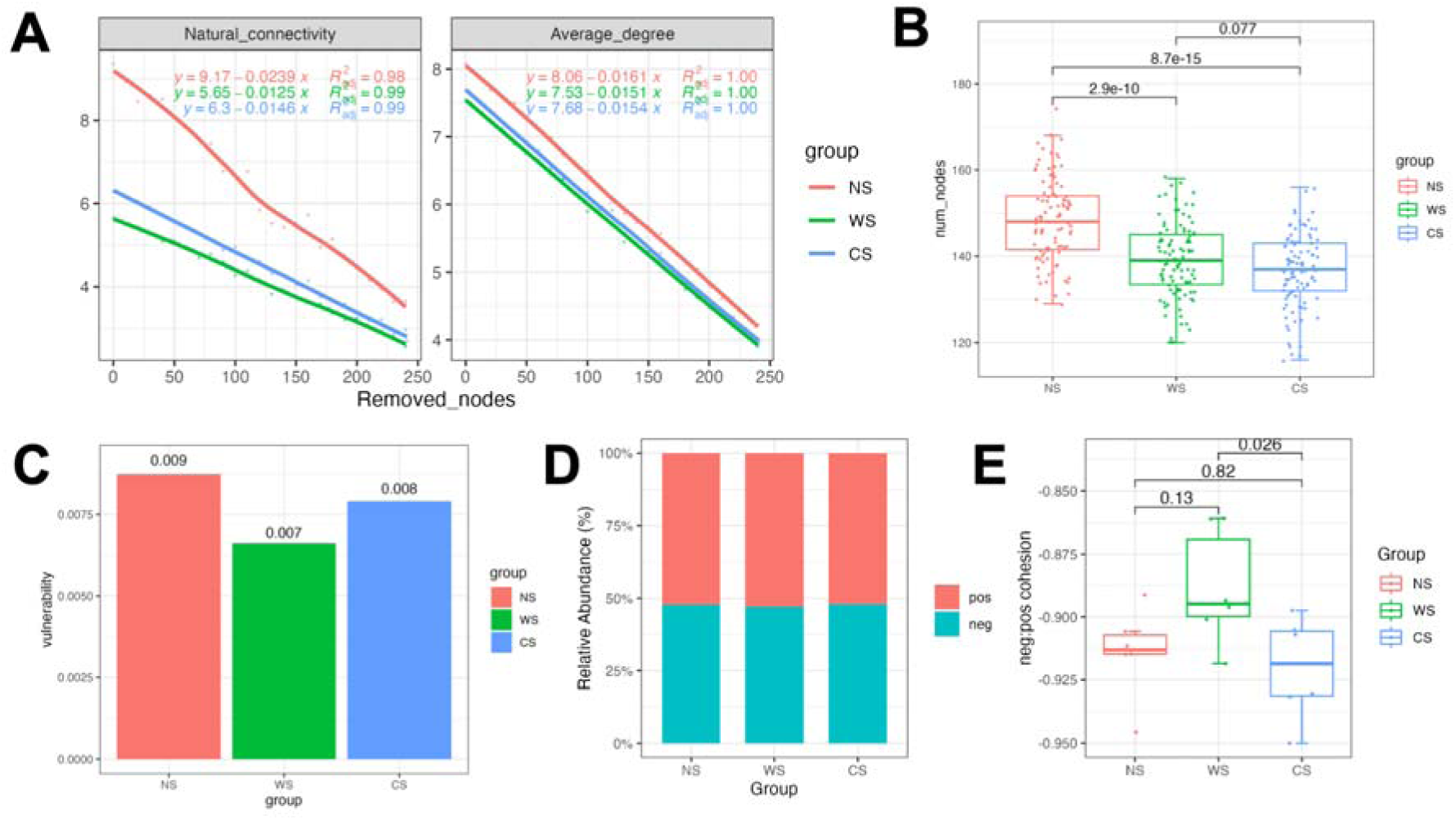
Network robustness and cohesion metrics. (A) Changes in natural connectivity and average degree during node removal simulation in three subnetworks. (B) Box plots comparing robustness scores of the three subnetworks. (C) Calculated vulnerability indices for each subnetwork. (D) Bar plots showing the relative proportions of positive and negative cohesion in the three subnetworks. (E) Box plots comparing the negative-to-positive cohesion ratios (neg: pos) among the subnetworks.

**Figure S6.**
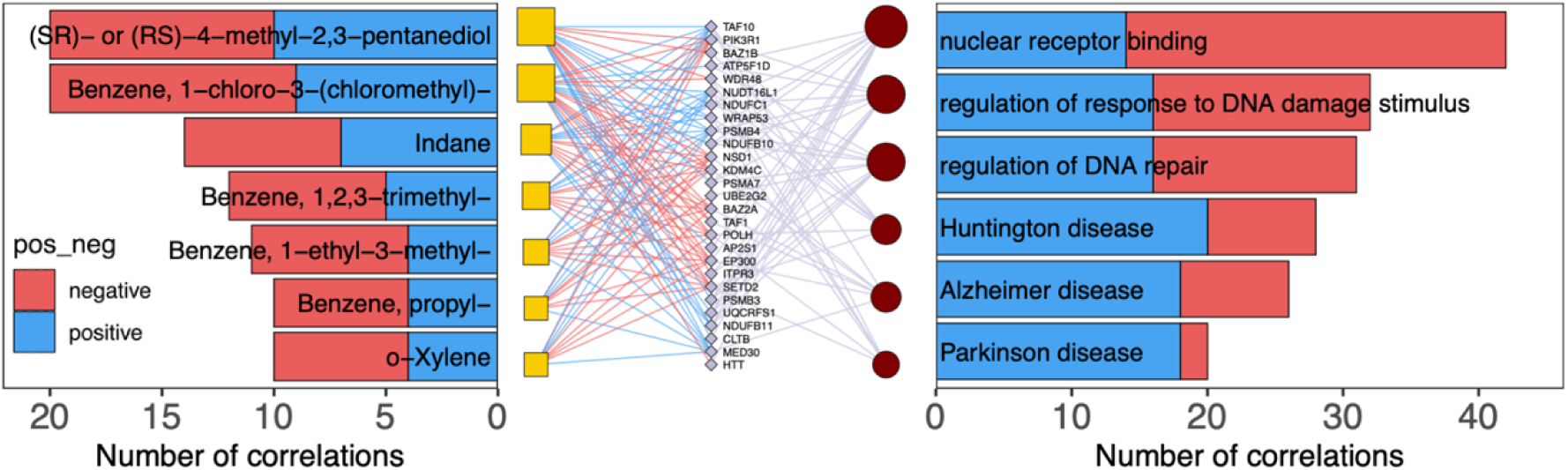
Correlation networks between the exposome and host transcriptome. Network representations of significantly correlated genes and enriched pathways for chemical exposures. Bar charts on either side indicate the number of positively and negatively correlated connections for each exposure.

